# Mechanosensitive Osteogenesis on Native Cellulose Scaffolds for Bone Tissue Engineering

**DOI:** 10.1101/2021.05.26.444470

**Authors:** Maxime Leblanc Latour, Andrew E. Pelling

## Abstract

In recent years, plant-derived cellulosic biomaterials have become a popular way to create scaffolds for a variety of tissue engineering applications. Moreover, such scaffolds possess similar physical properties (porosity, stiffness) that resemble bone tissues and have been explored as potential biomaterials for tissue engineering applications. Here, plant-derived cellulose scaffolds were seeded with MC3T3-E1 pre-osteoblast cells. Moreover, to assess the potential of these biomaterials, we also applied cyclic hydrostatic pressure (HP) to the cells and scaffolds over time to mimic a bone-like environment more closely. After one week of proliferation, cell-seeded scaffolds were exposed to HP up to 270 KPa at a frequency of 1Hz, once per day, for up to two weeks. Scaffolds were incubated in osteogenic inducing media (OM) or regular culture media (CM). The effect of cyclic HP combined with OM on cell-seeded scaffolds resulted in an increase of differentiated cells. This corresponded to an upregulation of alkaline phosphatase activity and scaffold mineralization. Importantly, the results reveal that well known mechanosensitive pathways cells which regulate osteogenesis appear to remain functional even on novel plant-derived cellulosic biomaterials.

## Introduction

Large defects created by either injury or disease may require graft placement to avoid non-union or malunion of the bone tissue (Andrzejowski and Giannoudis, 2019). Grafts can be derived directly from the patient (autologous grafts), and is considered the “gold standard” in regenerative orthopedics (Campana et al., 2014; Parikh, 2002; Sakkas et al., 2017; Wang and Yeung, 2017). However, limited size grafts, donor site morbidity and infections, cost and post-operative pain at both donor and receiver site has led to the development of alternative approaches (Parikh, 2002; Wang and Yeung, 2017): cadaver donors (allograft), animal sources (xenograft), or artificially derived (alloplastic). Such alternatives all have their own benefits and drawbacks, the later however provides a potential alternative with lower risk of transmitted diseases and infections, as well as overcoming the size limitation barrier (Parikh, 2002; Wang and Yeung, 2017). Alloplastic grafts are also considered a more ethical alternative to allografts and xenografts (Fernández et al., 2015). Physical properties are key parameters for graft development, such as pore size, interconnectivity and elasticity (Campana et al., 2014; Gao et al., 2019; Nukavarapu et al., 2015).

A spectrum of forces act on different areas the skeletal system. For instance, the pressure found in the femur head in human adults can reach 5 MPa during normal locomotion, and can reach up to 18 MPa for other activities (Morrell et al., 2005). On a microscopic level, these forces are transmitted to the osteocytes through Wnt/β-catenin mechano-sensing pathways in the lacuna-canaliculi network (Bonewald and Johnson, 2008). Force-regulated mechanisms lead to formation and removal of bone tissue through bone remodeling processes (Bonewald and Johnson, 2008) and the pressure inside the lacuna-canaliculi network is around 280 kPa (Zhang et al., 1998). Bioreactors have also been developed to apply stresses to replicate the native bone environment via uniaxial compression, tension, shear-stress, etc. (Brunelli et al., 2019; Pörtner et al., 2005). Pressure modulating bioreactors using low-intensity pulsed ultrasound (LIPUS) were also developed to stimulate osteoblastic and chondrogenic differentiation (Aliabouzar et al., 2018, 2016; Katiyar et al., 2014; Minto et al., 2020; Osborn et al., 2019; Zhou et al., 2016). Finally, hydrostatic pressure (HP) stimulation on cultured cells has also been achieved by compressing the gas phase above an incompressible media (Gardinier et al., 2009; Henstock et al., 2013; Liu et al., 2010; Reinwald et al., 2015; Reinwald and El Haj, 2018; Stavenschi et al., 2018; Zhao et al., 2015). Three-dimensional (3D) culturing of the cells is also critical for better representing *in vivo* conditions. With a specific scaffold structure and appropriate applied mechanical stimuli one can potentially predict the performance of a biomaterial scaffold prior to *in vivo* animal studies.

A variety of biomaterials have been utilized to mimic bone tissues. These include hydroxyapatite, tricalcium phosphate, bioceramics or bioactive glass (Wang and Yeung, 2017). These materials are osteoconductive, can promote osteointegration and provide structural support at the implant site (Wang and Yeung, 2017). However, they show little osteogenic response (Wang and Yeung, 2017). Polymer biomaterials such as poly(glycolic acid) (PGA), poly(lactic acid) (PLA) and poly(ε-caprolactone) (PCL) are also biocompatible, possess tunable degradation rates and can be chemically modify to change the surface chemistry (Wei et al., 2020). However, *in vivo* degradation creates acidic bi-products which can lead to an inflammatory response and decrease the efficiency bone repair (Wei et al., 2020). Finally, naturally occurring polymers such as collagen, chitosan and silk are also common (Wei et al., 2020). However, due to their inherent mechanical properties and structural stability, these materials are often utilized as composites with additional polymers and coatings in BTE applications (Di Martino et al., 2005; Lee and Volpicelli, 2017; Wei et al., 2020). More recently, plant-derived decellularized cellulose scaffolds have been shown to be effective in BTE applications (Hickey and Pelling, 2019; Lee et al., 2019; Torgbo and Sukyai, 2018).

Previous studies by our group and others have shown that cellulose based scaffolds derived from plants can be used as tissue engineering scaffolds (Bilirgen et al., 2021; Cheng et al., 2020; Contessi Negrini et al., 2020; Dikici et al., 2019; Fontana et al., 2017; Gershlak et al., 2017; Harris et al., 2021; Hickey et al., 2018; Holmes et al., 2022; Lee et al., 2019; Modulevsky et al., 2016, 2014; Robbins et al., 2020; S. H et al., 2021; Toker et al., 2020; Wang et al., 2020; Zhu et al., 2021). These biomaterials are often sourced from plants with a microstructure that closely mimics the tissue to be replicated (Hickey et al., 2018). Successful experiments *in vitro* and *in vivo* showed that these biomaterials are biocompatible and support angiogenesis (Hickey et al., 2018; Lee et al., 2019; Modulevsky et al., 2016, 2014). Other groups have successfully differentiated human pluripotent stem cells into bone-like tissues within scaffolds derived from either decellularized mushrooms (Balasundari et al., 2012), carrot (Contessi Negrini et al., 2020), bamboo stem (S. H et al., 2021) or apple tissues (Lee et al., 2019). The in vivo performance of the apple-derived scaffolds were further examined by implanting disk-shaped scaffolds in rat cranial defects (Lee et al., 2019). The findings demonstrate partial bone regeneration within the implant, type 1 collagen deposition and blood vessel formation (Lee et al., 2019). However, to date, the mechanobiology of cells cultured on plant-derived scaffolds has not been examined. It remains poorly understood how mechanical signal transduction pathways are, or are not, impacted when cultured on plant-derived cellulosic biomaterials. Here, to further examine the potential role plant-based biomaterials can play in BTE applications, we explore how mechanical stimulation impacts the differentiation of pre-osteoblasts when cultured on plant cellulose scaffolds. In a custom-built bioreactor, we apply cyclic HP stimulation to differentiating osteoblasts and examine the regulation of key markers of osteogenesis and mineralization. The results reveal that application of HP, in combination with osteogenic inducing media, leads to enhanced differentiation for cell-seeded MC3T3-E1 native cellulose scaffolds, and no significant change in the Young’s modulus of the scaffolds. This work provides further evidence that plant-derived cellulose scaffolds support osteogenesis and have potential applications in BTE.

## Materials and Methods

### Scaffold fabrication

Samples were prepared following established protocols (Hickey et al., 2018; Modulevsky et al., 2016, 2014). MacIntosh apples were cut with a mandolin slicer to 1 mm-thick slices. A biopsy punch was used to create 5 mm-diameter disks in the hypanthium tissue. The disks were decellularized in a 0.1% sodium dodecyl sulfate solution (SDS, Fisher Scientific, Fair Lawn, NJ) for 48h. Then, the decellularized disks were washed in deionized water before incubation in 100 mM CaCl_2_ for 48h. The samples were sterilized with 70% ethanol and placed in a 96-well culture plate.

MC3T3-E1 Subclone 4 cells (ATCC® CRL-2593™, Manassas, VA) were cultured in Minimum Essential Medium (ThermoFisher, Waltham, MA), supplemented with 10% Fetal Bovine Serum (Hyclone Laboratories Inc., Logan, UT) and 1% Penicillin/Streptomycin (Hyclone Laboratories Inc). Scaffolds were immersed in culture media and incubated at 37°C, 5% CO_2_, for 30 min. Cells were suspended and a 30 μL drop containing 5 × 10^4^ cells, was pipetted on each scaffold. The cells were left to adhere for 2 hours before adding 200 μL of culture media. Culture media was then changed every 3-4 days for 1 week. Cell seeded scaffolds were then either incubated in osteogenic media (OM) by adding 50 μg/mL of ascorbic acid and 10 mM β-glycerophosphate to the culture media or incubated in culture media (CM) for 2 weeks, with or without the application of HP.

### Cyclic hydrostatic pressure stimulation

Cyclic HP was applied by modulating the pressure in the gas phase above the culture wells in a custom-build pressure chamber (Figure 1, A). The humidified incubator atmosphere was compressed using an air compressor. A Particle Photon microcontroller (Particle Industries, San Francisco, CA) was used to control the frequency of the applied pressure remotely via a custom-made cellphone application through the Blynk IoT platform (Blynk, New York, NY). Cyclic HP was applied 1 hour per day, for up to 2 weeks (Figure 1, B) at a frequency 1Hz, oscillating between 0 and 280 kPa with respect to ambient pressure. Pressure was monitored using a pressure transducer. Samples were removed from the pressure chamber after each cycle and kept at ambient pressure between the stimulation phases.

**Figure 1:**
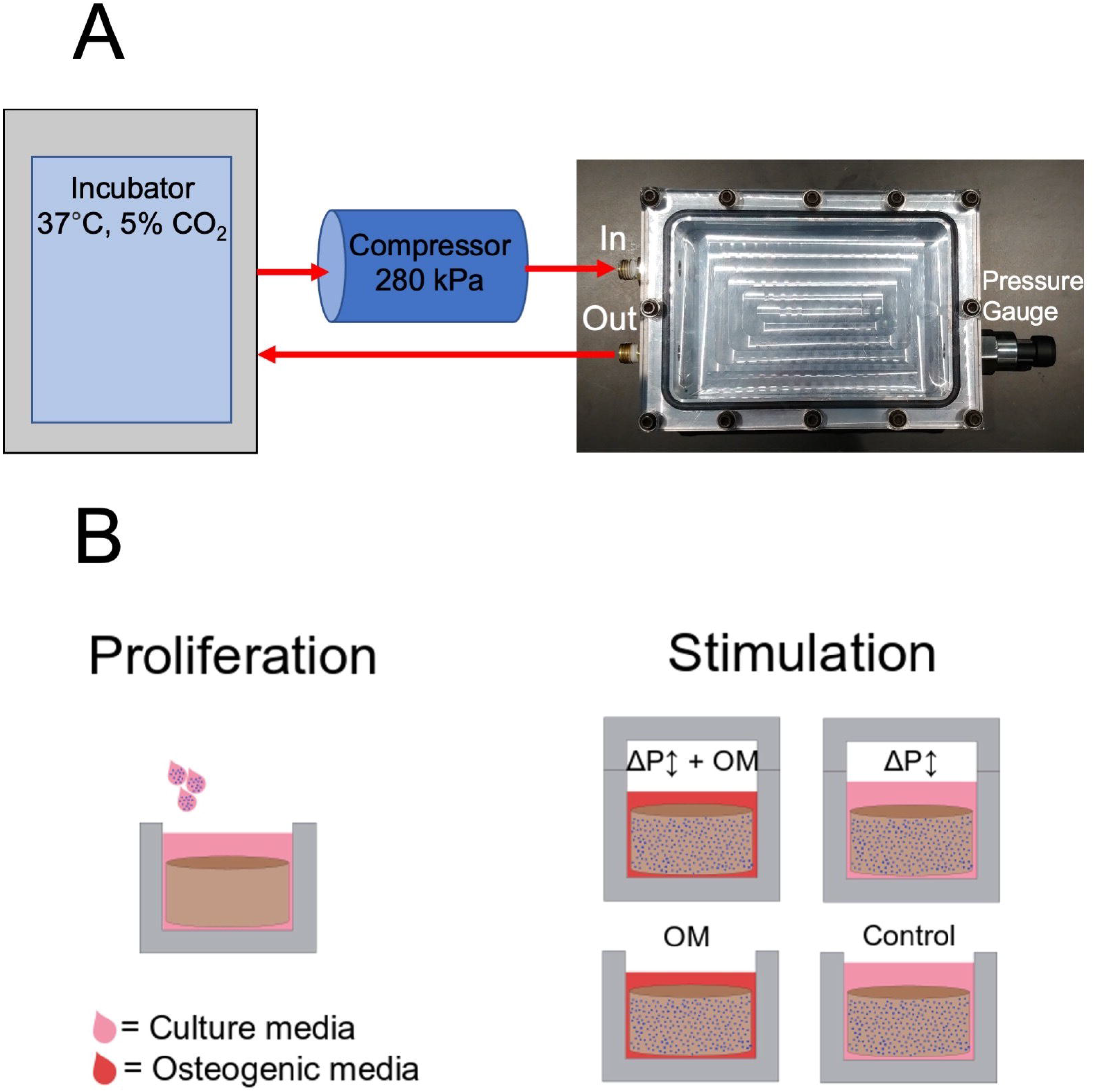
(A) Cyclic hydrostatic pressure device schematics. Hydrostatic pressure was applied by modulating the pressure in the gas phase above the culture wells in a custom-build pressure chamber. Air from incubator atmosphere was compressed using a compressor and injected in the pressure chamber using solenoid valves. (B) Experimental conditions. After 1 week of proliferation, cyclic hydrostatic pressure stimulation was applied during 1 hour per day, for up to 2 weeks at a frequency 1Hz, oscillating between 0 and 280 kPa with respect to ambient pressure. The samples were removed from the pressure chamber after each cycle and kept at ambient pressure between the stimulation phases.

Cell-seeded scaffolds were either stimulated with cyclic HP with and without the presence of OM, leading to four experimental conditions (Figure 1, B): Cyclic HP in regular culture media (CM-HP), cyclic HP in osteogenic culture media (OM-HP), non-stimulated in osteogenic media (OM-CTRL) and non-stimulated in regular culture media (CM-CTRL).

### Scaffold imaging

Scaffolds were washed with PBS and fixed with 10% neutral buffered formalin for 10 min. Scaffolds were washed with PBS and incubated in a 0.01% Congo Red staining solution (Sigma-Aldrich, St. Louis, MO) for 20 min at room temperature. Cell nuclei were stained with 1:1000 Hoechst (ThermoFisher, Waltham, MA) for 30 min. Samples were washed with PBS and stored in wash buffer solution (5% FBS in PBS). The cell-seeded scaffolds were imaged with a laser scanning confocal microscope (Nikon Ti-E A1-R) equipped with a 10X objective. Maximum intensity projections were used for cell counting with ImageJ software (Schindelin et al., 2012). Cells were counted on a 1.3 by 1.3 mm^2^ area (N=3 per experimental conditions with 3 randomly selected area per scaffold).

### Alkaline phosphatase activity assay

Alkaline phosphatase (ALP) activity in media was measured using an ALP assay kit (BioAssay Systems, Hayward, CA). Working solution was prepared with 5 mM magnesium acetate and 10 mM p-nitrophenyl phosphate (pNPP) in assay buffer, following manufacturer’s protocol. 150 μL of working solution was pipetted in 96-well plate. 200 μL of calibrator solution and 200 μL of dH_2_O were pipetted in separated well, in the same 96-well plate. At 1 week and 2 weeks, 20 μL of incubation media was pipetted into the working solution’s well. All wells were read at 405 nm for 10 minutes, every 30 seconds. ALP activity was calculated by taking the slope of the 405 nm readings vs time. Wells were read in triplicates (N=3 per experimental conditions).

### Alizarin red S staining and mineral deposit quantification

Samples were fixed with 10% neutral buffered formalin for 10 min, after 1 week or 2 weeks. Calcium quantification was performed using previously published protocol (Gregory et al., 2004). Samples were transferred to a 24-well plate and carefully washed with deionized water and incubated in 1 mL of 40mM (pH=4.1) alizarin red s (ARS, Sigma-Aldrich) solution for 20 minutes at room temperature, with light agitation. The samples were washed with deionized water and placed in 15 mL tubes filled with 10 mL dH_2_O. The tubes were placed on a rotary shaker at 120 rpm for 60 min and dH_2_O was replaced every 15 min. Thereafter, samples were incubated in 800 μL of 10% acetic acid on an orbital shaker at 60 rpm for 30 min. The eluted ARS/acetic acid solution was transferred to 1.5 mL centrifuge tubes. Tubes were centrifuged at 17 × 10^4^ g for 15 min. 500 μL of supernatants were transferred to new centrifuge tube and 200 μL of 10% ammonium hydroxide was added. Finally, 150 μL of the solution was pipetted into a 96-well plate and the absorption at 405nm was read using a plate reader. Wells were read in triplicates (N=3 per experimental conditions).

### Young’s modulus measurements

Young’s modulus measurements of the scaffolds were performed using a custom-built uniaxial compression apparatus, as previously described (Hickey et al., 2018). Briefly, scaffolds were mechanically compressed at a rate of 3 mm/min. The Young’s modulus of the scaffolds under the different experimental conditions were obtained by fitting the linear region of the stress-strain curve.

### Statistical analysis

Reported values are the average value ± standard error of the mean (SEM). Statistical significance was determined using one-way ANOVA and post hoc Tukey test. A value of *p* < 0.05 was considered to be statistically significant.

## Results

### Scaffold imaging and cell counting

The application of HP significantly increases the density of cells (Figure 2) after 1 week in OM compared to the static condition (p=10^−5^), but the increase was not significant after 2 weeks (p=0.07). Conversely, in CM a non-significant increase in the density of cells was observed with applied HP after 1 week (p=0.21) and 2 weeks (p=0.92). Importantly, we also observed a significant increase when incubated in OM compared to CM after 1 week of HP stimulation (p=0.02). After 2 weeks of HP stimulation, samples cultured in OM exhibited a similar density to samples cultured in CM (p=0.23). The results indicate that cell density increases more rapidly in the first week of stimulation in OM compared to CM media but that by two weeks the cell densities become equal. No significant difference was observed in the static cases after 1 or 2 weeks (p=0.99 in both cases).

**Figure 2:**
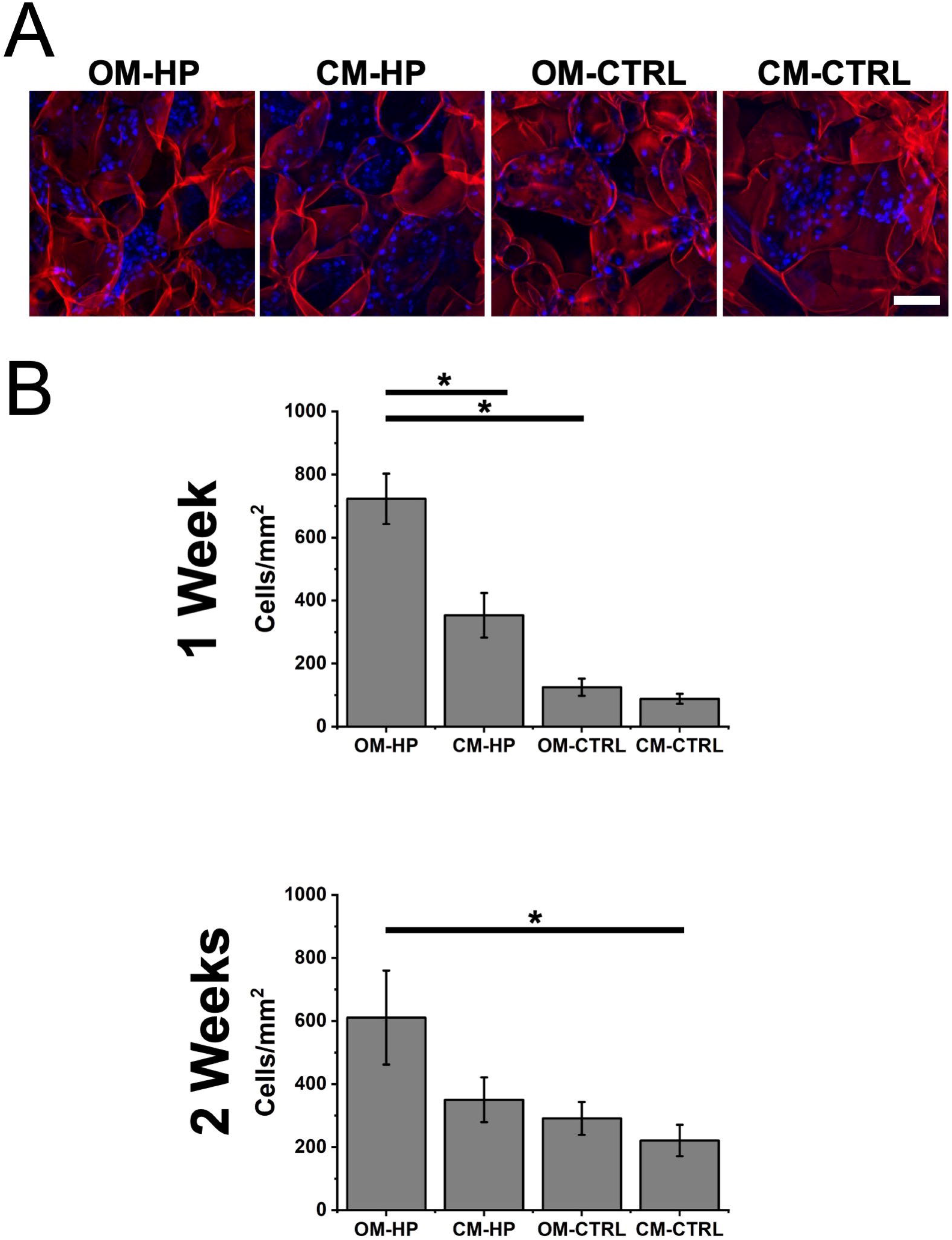
(A) Representative confocal laser scanning microscope image showing seeded cells scaffolds (scale bar = 100 μm – applies to all). The scaffolds were stained for cellulose (red) and for cell nuclei (blue). (B) Cellular density after 1 week or 2 weeks of stimulation. Statistical significance (* indicates p<0.05) was determined using a one-way ANOVA and Tukey post-hoc tests. Data are presented as means ± S.E.M. of three replicate samples per condition, with three areas per sample. OM-HP: Osteogenic Media – High Pressure; CM-HP: Culture Media – High Pressure; OM-CTRL: Osteogenic Media – Atmospheric Pressure; CM-CTRL: Culture Media – Atmospheric Pressure.

### Alkaline phosphatase activity assay

The stimulation with cyclic HP significantly increased the ALP activity (Figure 3) in scaffolds incubated in OM after 1 and 2 weeks compared to static condition (p=4×10^−8^ in both cases). A similar effect was observed in CM after 1 and 2 weeks (p=0.03 and p=5×10^−8^ respectively). However, the incubation in OM significantly increased ALP activity when HP is applied compared to incubation in CM, after 1 week (p<10^−8^) but was not significantly different after 2 weeks (p=0.99). Consistent with the cell density data the HP-driven increases in ALP activity are only observed during the first week and equalize by the second week of culture. In the absence of HP, the choice of incubation media did not significantly change the ALP activity after 1 or 2 weeks (p=0.25 and p=0.08 respectively).

**Figure 3:**
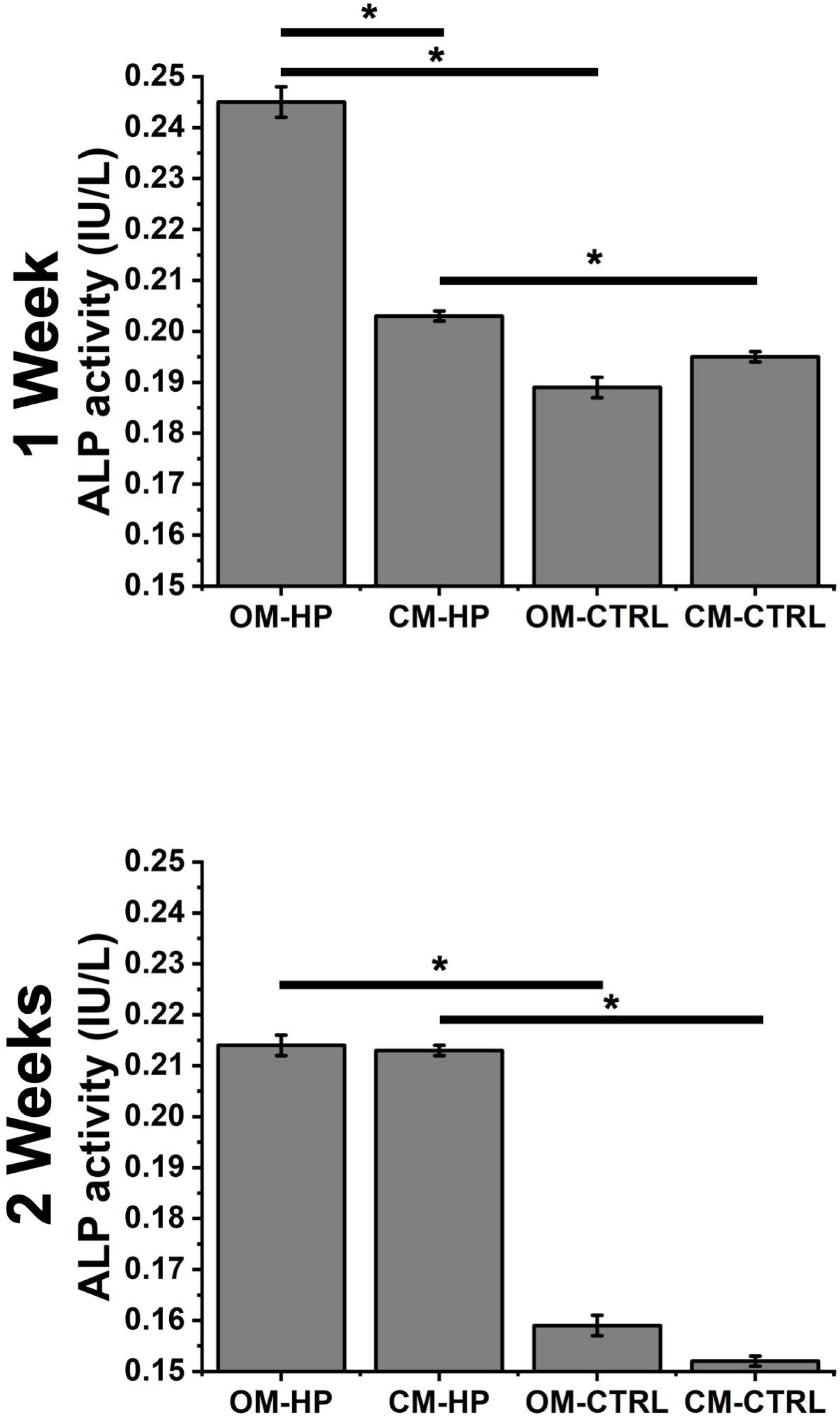
Alkaline phosphatase (ALP) activity after 1 week or 2 weeks of stimulation. Statistical significance (* indicates p<0.05) was determined using a one-way ANOVA and Tukey post-hoc tests. Data are presented as means ± S.E.M. of three replicate samples per condition. OM-HP: Osteogenic Media – High Pressure; CM-HP: Culture Media – High Pressure; OM-CTRL: Osteogenic Media – Atmospheric Pressure; CM-CTRL: Culture Media – Atmospheric Pressure.

### Alizarin red S staining and mineral deposit quantification

The application of cyclic HP significantly increased mineral deposition (Figure 4) for samples incubated in OM compared to static condition after 1 week and 2 weeks (p=2×10^−7^ and p=2×10^−8^ respectively). Similarly in samples cultured in CM, cyclic HP significantly increased mineral deposition after 1 week and 2 weeks (p=1×10^−6^ and p=2×10^−8^ respectively). Moreover, the incubation in OM significantly increased mineral deposition when HP is applied compared to incubation in CM, after 1 week (p=2×10^−4^) but was not significant after 2 weeks (p=0.99). These results are again consistent with the findings from assays of cell density and ALP activity. Under static conditions mineralization still occurred in OM as expected and was significantly increased compared to CM after 1 week (p=10^−3^) but was not significant after 2 weeks (p=0.75).

**Figure 4:**
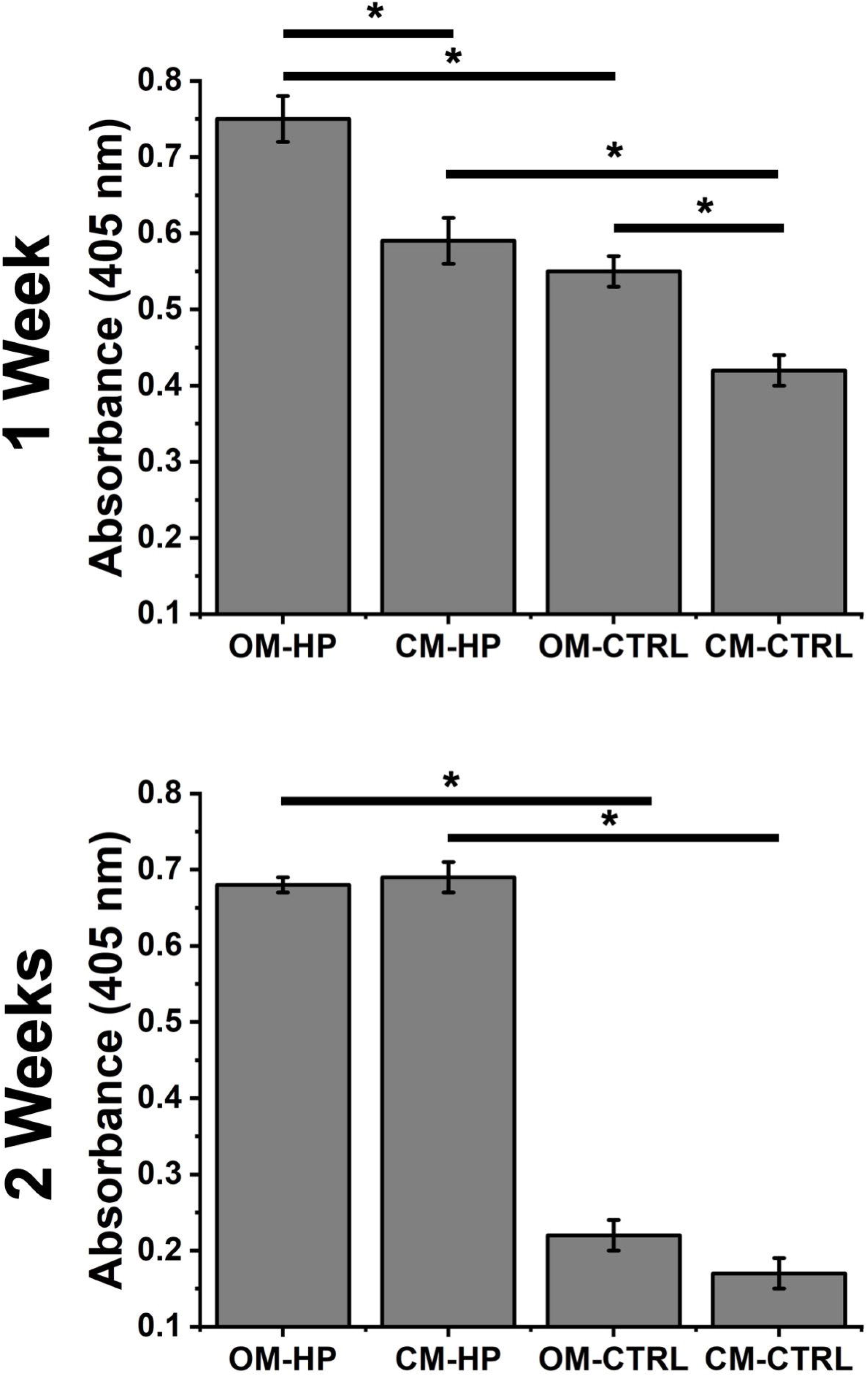
Mineral deposit quantification with Alizarin Red S (ARS) staining after 1 week or 2 weeks of stimulation. Statistical significance (* indicates p<0.05) was determined using a one-way ANOVA and Tukey post-hoc tests. Data are presented as means ± S.E.M. of three replicate samples per condition. OM-HP: Osteogenic Media – High Pressure; CM-HP: Culture Media – High Pressure; OM-CTRL: Osteogenic Media – Atmospheric Pressure; CM-CTRL: Culture Media – Atmospheric Pressure.

### Young’s modulus measurements

Scaffolds were assessed for change in Young’s modulus after stimulation (Figure 5). Data showed no significant changes between samples incubated in OM with applied HP (16.1 ± 2.1 kPa) and without applied HP (17.2 ± 3.2 kPa) after 1 week and 2 weeks (13.9 ± 0.8 kPa and 18.7 ± 0.7 kPa, respectively). Moreover, no significant change was observed in samples incubated in CM with HP or without HP, both after 1 week (14.2 ± 2.0 kPa and 13.9 ± 0.6 kPa, respectively) or 2 weeks (20.2 ± 2.3 kPa and 14.1 ± 4.7 kPa, respectively). Furthermore, no significant change was observed due to incubation in OM compared to CM, for samples under applied HP after 1 week (16.1 ± 2.1 kPa and 14.2 ± 2.0 kPa, respectively) or 2 weeks (13.9 ± 0.8 kPa and 20.2 ± 2.3 kPa, respectively). Similarly, no significant change was at atmospheric pressure comparing OM and CM at 1 week (17.2 ± 3.2 kPa and 14.2 ± 2.0 kPa, respectively) or after 2 weeks (18.7 ±.7 kPa and 14.1 ± 4.7 kPa, respectively).

**Figure 5:**
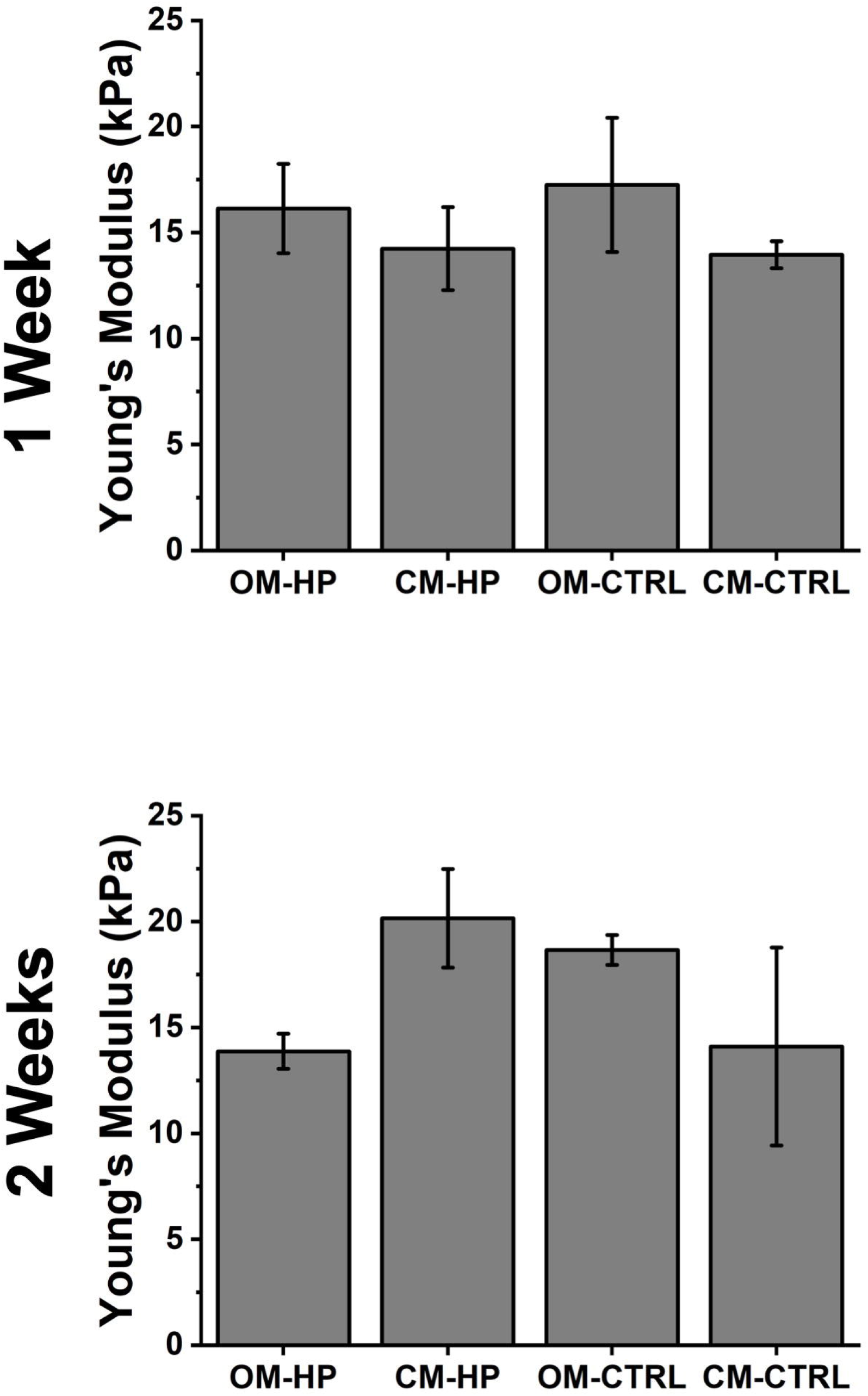
Young’s modulus of scaffolds after 1 week or 2 weeks of stimulation. No statistical significance was observed between any of the conditions. Data are presented as means ± S.E.M. of three replicate samples per condition. OM-HP: Osteogenic Media – High Pressure; CM-HP: Culture Media – High Pressure; OM-CTRL: Osteogenic Media – Atmospheric Pressure; CM-CTRL: Culture Media – Atmospheric Pressure.

## Discussion

Cells utilize a variety of mechanisms to sense and respond to a variety of mechanical stimuli (Vining and Mooney, 2017). Mechanical stimuli are known to affect cell differentiation, tissue regeneration, cytokines and protein expression and proliferation (Martino et al., 2018; Vining and Mooney, 2017). Bones are subjected to constant mechanical stresses and adapt through remodeling process (Bonewald and Johnson, 2008). *In vivo*, HP stimulates bone cells and impacts cell differentiation, maker expression and mineralisation (Henstock et al., 2013; Huang and Ogawa, 2012; Reinwald and El Haj, 2018). Plant-derived cellulose scaffolds are an emerging biomaterial in BTE (Lee et al., 2019), and therefore it is of interest to understand their performance under the mechanical conditions that are found *in vivo* (Gao et al., 2019; Nukavarapu et al., 2015). Cellulose biomaterials derived from plant tissues have shown promising results *in vitro* and *in vivo* for targeted tissue engineering (Hickey et al., 2018; Modulevsky et al., 2016, 2014) and have been used to host osteoblastic differentiation (Lee et al., 2019). In addition, apple-derived cellulose scaffolds exhibit similar morphological characteristics to trabecular bone and were previously used for BTE applications (Lee et al., 2019).

In this study, we replicated the mechanical stimuli present during human locomotion and measured the impact of differentiation markers in plant-derived cellulose scaffolds seeded with pre-osteoblast cells. External pressure was applied on the scaffolds in similar magnitude of the lacuna-canaliculi network with a frequency mimicking human locomotion (Henstock et al., 2013; Zhang et al., 1998). After proliferation, our scaffolds were either cultured in standard culture media, or in osteogenic-inducing differentiation media, with or without high pressure stimulation. Other groups utilizing similar cell lines (Gardinier et al., 2014, 2009), bone marrow skeletal stem cells (Reinwald and El Haj, 2018; Stavenschi et al., 2018; Zhao et al., 2015) or ex-vivo chick femur (Henstock et al., 2013) have also studied the effects if cyclic HP on either 2D surfaces, biomaterial meshes or ex vivo bones. In general, our results show a sensitivity of HP during the early differentiation of the cells during the first week of stimulation. This sensitivity largely subsides during the second week of culture.

Cell counting revealed that the application of HP enhances MC3T3-E1 proliferation when cultured in OM or CM. Consistent with our work, results from previous studies in which metabolic activity has also been shown to be upregulated by mechanical stimulation in comparison to non-stimulated samples (Reinwald and El Haj, 2018). Moreover, it was shown that the application of HP accelerates cell proliferation through upregulated cell cycle initiation (Zhao et al., 2015). Similarly, reports have shown that physical stimulation of MC3T3-E1 cells induced expression of paracrine factors that leads enhancement of cell proliferation (Stavenschi et al., 2018). Additionally, other type of cyclic pressure stimulation, such as ultrasound stimulation are known to increase the proliferation of MC3T3-E1 cells (Katiyar et al., 2014) and human mesenchymal stem cells (hMSC) (Aliabouzar et al., 2018, 2016; Osborn et al., 2019; Zhou et al., 2016). Importantly, when cultured in OM, cell density increases more rapidly with HP compared to CM during the first week of culture, but the cell densities equalize by the end of the second week. This data is consistent with previous reports of a time-dependent increase in cell number when cultured in OM (Quarles et al., 1992). Similarly, no significant difference between incubation of MC3T3-E1 cells in similar OM after 2 weeks (Hong et al., 2010). Our findings corroborate these studies and further suggest that the application of HP influences the replication rate at early stages of stimulation for samples cultured in OM.

ALP is an important enzyme expressed in the early stages of osteoblastic differentiation (Golub and Boesze-Battaglia, 2007). Our results indicate that the application of cyclic HP significantly increases ALP activity, compared to the static case. These findings are consistent with other studies on more conventional scaffolds (Reinwald and El Haj, 2018). For example, a significant increase in ALP activity was also reported after the incubation of scaffolds in osteogenic-inducing differentiation media, similarly to reports on 2D culture systems (Hong et al., 2010; Quarles et al., 1992). Other reports showed that ultrasound stimulation increased ALP activity in osteoblast-differentiated hMSC on polyethylene glycol diacrylate (PEGDA) (Zhou et al., 2016) and polylactic acid (PLA) (Osborn et al., 2019) scaffolds, after 3 weeks. The application of HP significantly increased the mineral content in the scaffolds after 1 week and 2 weeks of stimulation, in both types of incubation media. Other groups have shown that a cyclic 300 kPa pressure at 2 Hz frequency on human BMSCs promoted significant mineral deposition (Stavenschi et al., 2018). The increase in mineral deposition also noted in ex vivo bone samples, with similar HP force application (Henstock et al., 2013). Furthermore, the incubation in OM increased the mineral content in the scaffolds, which is consistent with other studies (Hong et al., 2010; Quarles et al., 1992). The increased in mineral content was also observed by other groups for osteoblast-differentiated hMSC stimulated with ultrasound, for up to 13% in extracellular calcium deposition (Osborn et al., 2019; Zhou et al., 2016). Along with ALP expression, mineral content expression further confirms the ongoing differentiation of MC3T3-E1 onto osteoblast, either by applied HP, chemically (induction in OM) or a combination of both.

Although the measured mechanical properties of the scaffolds were consistent with previous studies (Aliabouzar et al., 2018; Burger et al., 2020; Contessi Negrini et al., 2020; Harris et al., 2021; Kim et al., 2018; Maharjan et al., 2021a; Osborn et al., 2019; Zaborowska et al., 2010; Zhou et al., 2016) analysis revealed that the application of cyclic hydrostatic pressure did not impact their elastic properties. Results showed that Young’s moduli of the scaffolds (13.9-20.2 kPa, Figure 5) are similar to previous reports using apple-derived scaffolds, at 4.17 ± 0.17 kPa (Contessi Negrini et al., 2020). Although our values are slightly higher, the small discrepancy could be due to the type of cultivar used, *Golden delicious* vs *McIntosh*. Regardless, our results fall into a broad range of reported values by other groups using other types of decellularized plant cellulose as scaffolds (Harris et al., 2021). Similarly, our results are comparable to other cellulose-based biomaterials for BTE: Scaffolds from regenerated cellulose fibers and chitosan, with cultured MC3T3-E1 cells have a reported Young’s modulus of up to 0.017 kPa (Maharjan et al., 2021b). Cellulose-silk hydrogel was reported to be ranging from 53-72 kPa (Burger et al., 2020). Moreover, cellulose-based scaffolds with a higher Young’s modulus (in MPa) were also reported (Kim et al., 2018; Zaborowska et al., 2010). Other groups also reported Young’s moduli of synthetic scaffolds for BTE, in combination with ultrasound stimulation: PEG-DA, ranging from 0.84 MPa to 2.63 MPa (Aliabouzar et al., 2018; Zhou et al., 2016) or PLA, 2.15 MPa (Osborn et al., 2019). One should note that the reported Young’s moduli are lower than cortical and trabecular bone (15-20 GPa; 0.1-2 GPa, respectively) (Bose et al., 2012). Thus, it may be preferrable to limit potential *in vivo* applications to non-load bearing BTE.

## Conclusion

Plant-derived scaffolds have recently been demonstrated as an interesting alternative to autografts, xenografts and synthetics implants (Bilirgen et al., 2021; Cheng et al., 2020; Contessi Negrini et al., 2020; Dikici et al., 2019; Fontana et al., 2017; Gershlak et al., 2017; Harris et al., 2021; Hickey et al., 2018; Holmes et al., 2022; Lee et al., 2019; Modulevsky et al., 2016, 2014; Robbins et al., 2020; S. H et al., 2021; Toker et al., 2020; Wang et al., 2020; Zhu et al., 2021). However, it has remined poorly understood how mechanosensitive pathways in bone precursor cells are impacted by being cultured in plant-derived biomaterials. Here, mechanical pressur-driven stimulation has allowed us to characterize the mechanosensitive behaviors of bone precursor cells on these novel scaffolds. The results reveal that application of cyclic HP, in combination with OM, leads to an increase in the number of cells, ALP activity and mineralization over time, as compared to non-stimulated, static experiments. Results showed that the elastic properties of the scaffolds were not affected by the application of cyclic HP, nor the type of incubation media. Importantly, this work provides evidence that bone-precursor cells possess intact mechanosensing and mechanotransduction pathways when cultured on novel plant-derived scaffolds in a mechanically active environment. These results combined with past *in vitro* and *in vivo* studies using apple-derived scaffold biomaterials demonstrate their potential for some BTE applications.

## Acknowledgements

This work was supported by a Discovery Grant from the Natural Sciences and Engineering Research Council of Canada (NSERC) and a grant from the Li Ka Shing Foundation.

